# Identification of efflux substrates using a riboswitch-based reporter in *Pseudomonas aeruginosa*

**DOI:** 10.1101/2023.02.27.530370

**Authors:** Verónica Urdaneta-Páez, Randy Hamchand, Karen Anthony, Jason Crawford, Alan G. Sutherland, Barbara I. Kazmierczak

**Author notes:** Address correspondence to Barbara I. Kazmierczak.

## Abstract

*Pseudomonas aeruginosa* is intrinsically resistant to many classes of antibiotics, reflecting the restrictive nature of its outer membrane and the action of its numerous efflux systems. However, the dynamics of compound uptake, retention and efflux in this bacterium remain incompletely understood. Here, we exploited the sensor capabilities of a Z-nucleotide sensing riboswitch to create an experimental system able to identify physicochemical and structural properties of compounds that permeate the bacterial cell, avoid efflux, and perturb the folate cycle or *de novo* purine synthesis. In a first step, a collection of structurally diverse compounds enriched in antifolate drugs was screened for ZTP riboswitch reporter activity in efflux-deficient *P. aeruginosa*, allowing us to identify compounds that entered the cell and disrupted the folate pathway. These initial hits were then rescreened using isogenic efflux-proficient bacteria, allowing us to separate efflux substrates from efflux avoiders. We confirmed this categorization by measuring intracellular levels of select compounds in the efflux-deficient and - proficient strain using high resolution LC-MS. This simple yet powerful method, optimized for high throughput screening, enables the discovery of numerous permeable compounds that avoid efflux and paves the way for further refinement of the physicochemical and structural rules governing efflux in this multi-drug resistant Gram-negative pathogen.

**Importance:** Treatment of *Pseudomonas aeruginosa* infections has become increasingly challenging. The development of novel antibiotics against this multi-drug resistant bacterium is a priority, but many drug candidates never achieve effective concentrations in the bacterial cell due due to its highly restrictive outer membrane and the action of multiple efflux pumps. Here, we develop a robust and simple reporter system in *P. aeruginosa* to screen chemical libraries and identify compounds that either enter the cell and remain inside, or enter the cell and are exported by efflux systems. This approach enables developing rules of compound uptake and retention in *P. aeruginosa* that will lead to more rational design of novel antibiotics.

## Introduction

*Pseudomonas aeruginosa* is a ubiquitous Gram-negative bacterium capable of colonizing diverse environments such as the rhizosphere, plants, insects, and mammals (1-3). It is an opportunistic pathogen of clinical relevance that has been associated with hospital-acquired infections and ventilator-associated pneumonia (4). *P. aeruginosa* also causes serious infections in immunocompromised individuals and patients with wounds and burns (5, 6) and is the prevalent pathogen causing chronic lung infections in cystic fibrosis patients (7).

*P. aeruginosa* exhibits intrinsic and extrinsic resistance to a broad range of antimicrobial compounds, making the treatment of infections a challenge (8). Limited cell permeability, efflux systems, and the production of antibiotic-inactivating enzymes all contribute to intrinsic resistance (9). *P. aeruginosa*’s outer membrane contains specific and small channel proteins for the uptake of nutrients, rather than non-specific porins that allow for diffusion of larger molecules. This restricted permeability limits the entry of noxious compounds, including antibiotics (9-12). *P. aeruginosa* also expresses twelve resistance nodulation division (RND) efflux systems (11, 13, 14), five of which (MexAB-OprM, MexXY-OprM, MexCD-OprJ, MexEF-OprN and MexJK-OprM) can export diverse types of antibiotics (14, 15). Lastly, *P. aeruginosa* expresses enzymes that break down or modify antibiotics, including β-lactamases and aminoglycoside-inactivating enzymes (16, 17). *P. aeruginosa* can also acquire resistance during antibiotic treatment. This generally occurs either by horizontal gene transfer or by selection of beneficial mutations, such as those promoting the overexpression of efflux systems and antibiotic-inactivating enzymes, or the structural inactivation of porins (9, 11, 18, 19).

Many antibiotic discovery efforts rely on a non-specific whole-cell approach that begins with the selection of compounds that inhibit bacterial growth (20-23). Even if these methods provide a list of promising candidates, the elucidation of the antibacterial mode of action is difficult and often unfruitful (21, 22, 24, 25). Given the critical contribution to antibiotic resistance conferred by limited outer membrane permeability and the action of efflux pumps in *P. aeruginosa* (8, 26-29), we have developed an alternative screening approach to address the poorly understood dynamics of compound uptake, retention, and efflux in this bacterium. Our strategy encompasses two key elements. The first is the use of riboswitch-based reporters designed to detect more subtle and specific metabolic perturbations in actively growing cells, rather than growth inhibition or cell death. The second is the use, in parallel, of isogenic efflux-proficient and efflux-deficient (Δ*mexAB-oprM nfxB* Δ*mexCD-oprJ* Δ*mexJKL* Δ*mexXY* Δ*opmH*362 Δ*mexEF-oprN*) strains for our screen (Figure 1).

**Figure 1.**
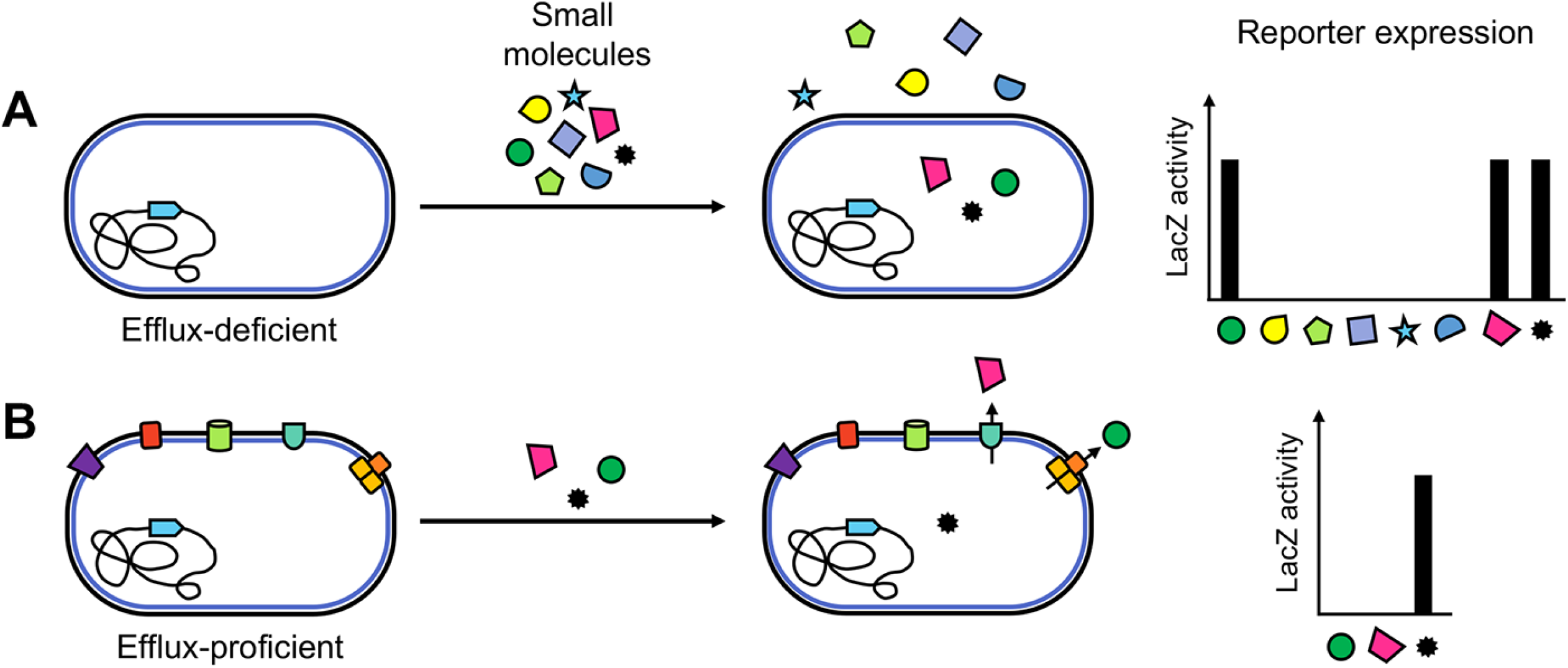
Compound screen outline using the ZTP-*lacZ* reporter. **A**. An efflux-deficient strain was employed to screen a small molecule library. Increased reporter expression identified compounds that enter the cell and inhibit the targeted pathway. **B**. Compounds identified in the first screen (A) were tested against an efflux-proficient strain. Some compounds now exhibited decreased or absent reporter expression, suggesting that they are efflux substrates, while for others reporter expression remained unchanged, suggesting that they are not subject to efflux.

Riboswitches are bacterial regulatory RNA elements often found in the 5’ untranslated region of messenger RNAs that control downstream gene expression as a function of binding metabolites, coenzymes, signaling molecules, or inorganic ion ligands (30). These natural biosensors allow bacteria to appropriately regulate gene expression in the setting of fluctuating concentrations of metabolites, metals, or salts. Riboswitch-based reporters have been used to characterize novel antibacterial drugs and targets (31, 32), to identify riboswitch ligands (33, 34), to describe biosynthetic pathways (35), and to screen for high-yield vitamin producing strains (36). To date, nearly 40 classes of riboswitches have been characterized, many of them monitoring essential metabolic pathways and controlling expression of virulence factors (37).

The ZTP riboswitch senses metabolic flux through the folate cycle and the *de novo* purine synthesis pathway (38). The folate pathway is constituted by six enzymes that convert guanosine triphosphate (GTP) to tetrahydrofolate (THF) (39). THF then serves as donor of one-carbon units in metabolic pathways that synthesize methionine, glycine, thymine, and purines, which are essential components of nucleic acids and proteins (40-42). The ZTP riboswitch tightly regulates these pathways by binding either ZTP (5-aminoimidazole-4-carboxamide riboside 5’-triphosphate) or its precursor ZMP (5-aminoimidazole-4-carboxamide ribonucleotide) and activating the expression of the enzymes from these pathways (38).

Here, we describe the implementation in *P. aeruginosa* of a riboswitch reporter system which couples the ZTP riboswitch sequence from *Pectobacterium carovotorum* (described in (38)) to *the lacZ* reporter gene. The reporter construct was introduced into isogenic efflux-proficient and efflux-deficient strains of *P. aeruginosa* and screened against a focused library of compounds enriched in antifolate drugs. By using an efflux-deficient strain, we identified all compounds that were cell permeable and inhibited enzymes in the folate pathway. We then re-tested these compounds against the efflux-proficient strain, allowing us to identify compounds that are likely substrates for efflux (Figure 1). The initial use of an efflux-deficient strain allowed the detection of active, cell-permeable compounds that would have been missed by using an efflux-proficient strain alone. These observations demonstrate the potential of our approach for establishing rules of compound uptake and retention in Gram-negative bacteria like *P. aeruginosa*.

## Results

### The *Pectobacterium carovotorum* ZTP riboswitch reporter responds to folate cycle inhibition in *Pseudomonas aeruginosa*

The ZTP riboswitch regulates expression of genes involved in the folate cycle and *de novo* purine synthesis pathway (38). To detect alterations of these pathways in *P. aeruginosa*, we developed a reporter system consisting of a translational fusion between the ZTP riboswitch sequence from *Pectobacterium carovotorum* and the *lacZ* reporter gene. This reporter construct was integrated into the *attB* site of the *P. aeruginosa* chromosome. The reporter fusion was expressed from the P*exoT* promoter (Figure 2A), which controls the expression of the type three secretion system (T3SS) effector ExoT (43). P*exoT* is activated by the AraC/XylS-type transcriptional regulator ExsA (44). The ExsA anti-activator ExsD (45, 46) was deleted in all reporter strains to allow for constitutive expression from the P*exoT* promoter (Table 1). Strains in which *exsA* was deleted, as well as strains harboring the M4 mutant allele of the ZTP riboswitch reporter (defective for ligand binding (38)), served as negative controls (Figure 2A, Table 1). Importantly, the M4 variant riboswitch, which no longer binds ZTP, allows compounds that directly bind and activate the substrate recognition structure of the riboswitch to be differentiated from compounds that disrupt the folate cycle.

**Figure 2.**
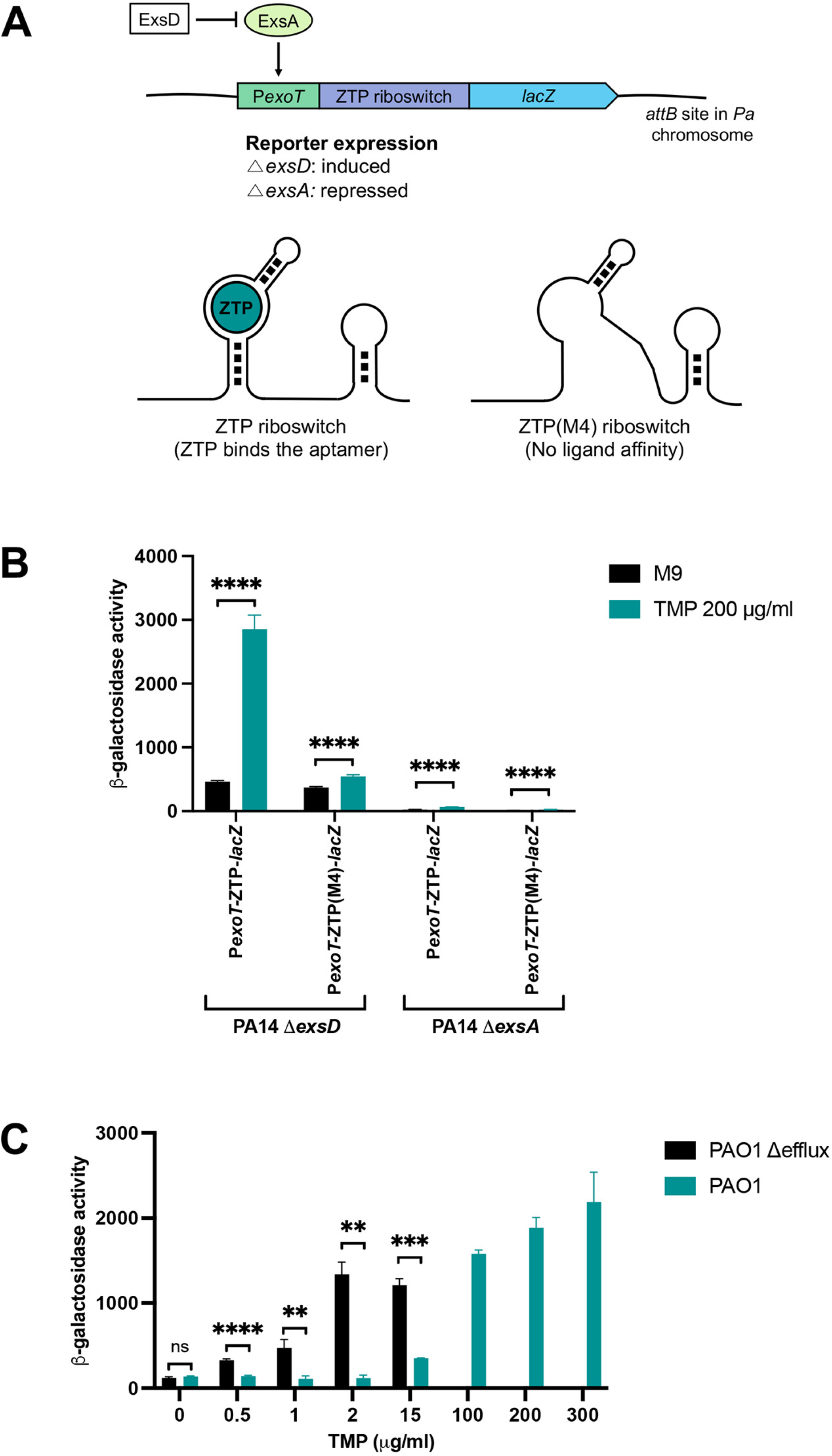
The ZTP riboswitch reporter is functional in *Pseudomonas aeruginosa*. **A**. The riboswitch reporter consists of the *lacZ* gene fused downstream from the *Pectobacterium carotovorum* ZTP riboswitch, with expression from the ExsA-regulated *exoT* promoter. The reporter is constitutively expressed in the absence of ExsD. Expression of genes downstream of the riboswitch requires ligand (ZTP) binding, and alteration of the aptamer structure by mutation (ZTP(M4) riboswitch) results in loss of ligand affinity. **B**. β-Galactosidase activity of translational P*exoT*-ZTP-*lacZ* and P*exoT*(M4)-ZTP-*lacZ* fusions in Δ*exsD* and Δ*exsA* backgrounds growing in M9 + 1% CAA (black histograms) and M9 + 1% CAA with 200 μg/mL TMP (teal histograms). Bars show means ± standard deviations from 3 independent experiments. Unpaired two-tailed t-tests were used to compare the β-galactosidase values of cultures grown in M9 + 1% CAA and cultures grown in M9 + 1% CAA with TMP. P < 0.0001, ****. **C**. β-galactosidase activity of the P*exoT*-ZTP-*lacZ* reporter in efflux-deficient (black histograms) and - proficient (teal histograms) bacteria grown in M9 + 1% CAA and exposed to TMP. Bars show means ± standard deviations from 3 independent experiments. Unpaired two-tailed t-tests were used to compare the efflux-deficient and -proficient strains at varying concentrations of TMP. P < 0.01, **; P < 0.001, ***; P < 0.0001, ****; not significant, ns.

**Table 1.**
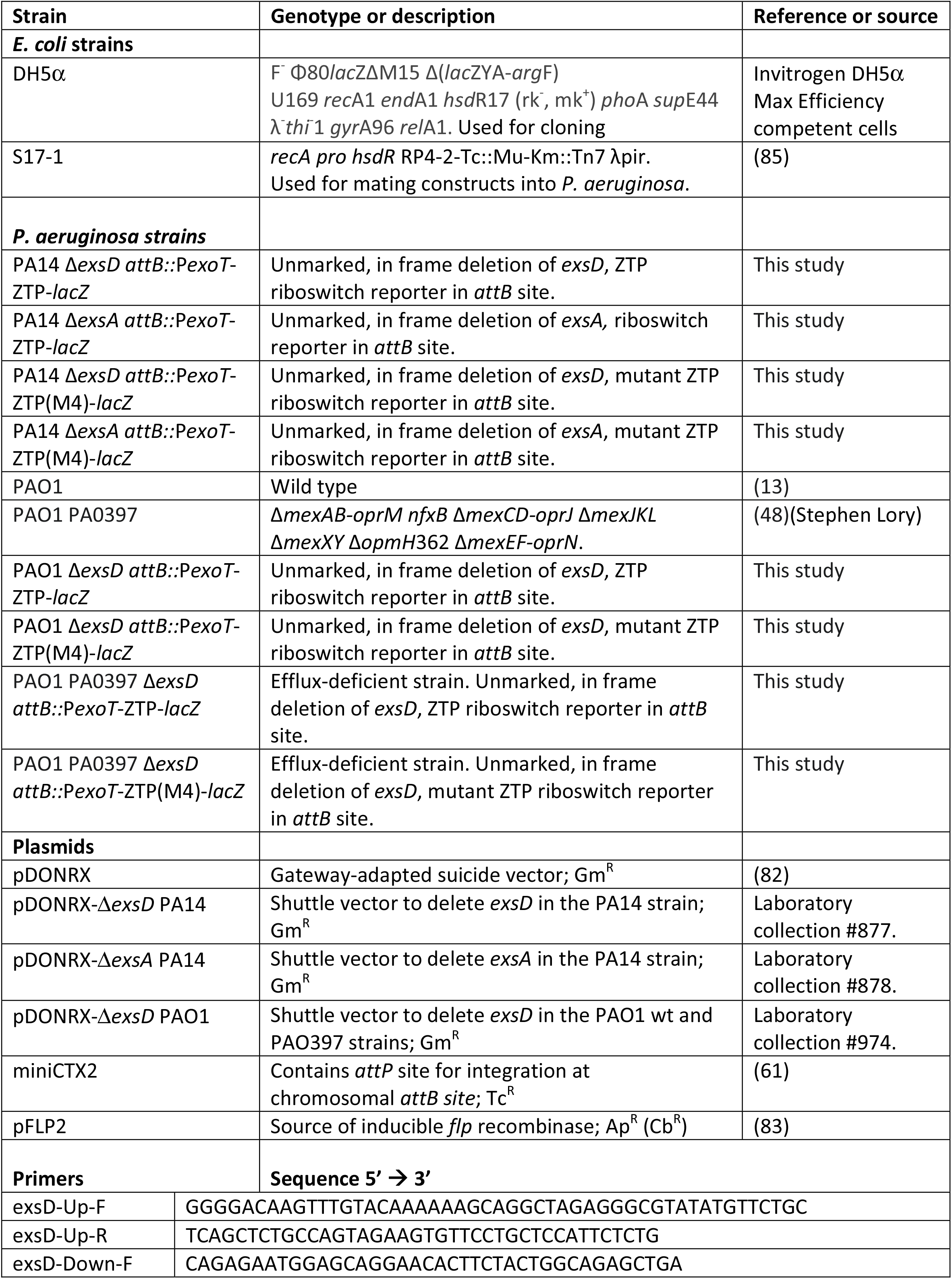
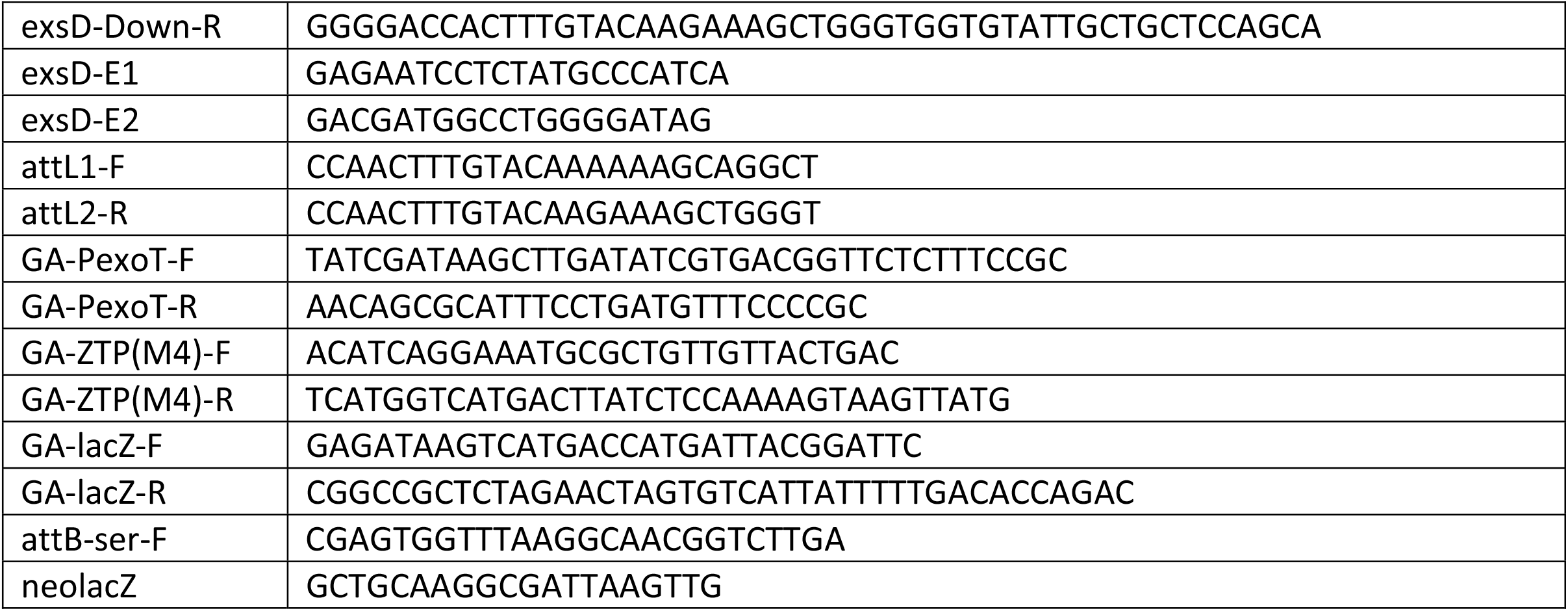
Bacterial strains, plasmids and primers used in this study.

To examine the activity of the riboswitch reporter constructs, bacteria were exposed for 3 hours to subinhibitory concentrations of trimethoprim (TMP, 200 μg/mL) before measuring β-galactosidase activity (Figure 2B). TMP is a dihydrofolate reductase (DHFR) inhibitor (47) that disrupts the folate pathway in bacteria, which results in increasing concentrations of the purine biosynthesis intermediate ZTP within the bacterial cell (38). As observed in figure 2B, treatment with TMP significantly increased reporter activity in the PA14 Δ*exsD attB::*P*exoT*-ZTP-*lacZ* strain, but failed to do so in the strain carrying the mutant ZTP riboswitch (PA14 Δ*exsD attB::*P*exoT*-ZTP(M4)-*lacZ*). Reporter activity was not detected in PA14 Δ*exsA* strains following treatment with TMP, as predicted by the ExsA dependence of the P*exoT* promoter (44). These results demonstrate that the riboswitch reporter construct is responsive to the antifolate drug TMP when tested in *P. aeruginosa*, and that the observed response is strictly dependent on ligand (ZTP) binding to the riboswitch aptamer region.

### Absence of active efflux significantly increases ZTP riboswitch reporter sensitivity

To assess ZTP reporter behavior in an efflux-deficient background, we integrated the ZTP reporter into PA0397 (PAO1 Δ*exsD* Δ*mexAB-oprM nfxB* Δ*mexCD-oprJ* Δ*mexJKL* Δ*mexXY* Δ*opmH*362 Δ*mexEF-oprN*)(48). This mutant carries deletions in multiple efflux systems previously shown to promote antibiotic export (14, 15, 28, 49, 50) and exhibits an increased sensitivity to trimethoprim (TMP), with a minimal inhibitory concentration (MIC) of 15 μg/mL as compared to the 450 μg/mL MIC observed for the PAO1 wt strain. We used lower concentrations of TMP (0.5 to 15 μg/mL) to test reporter activity in the efflux-deficient background (Figure 2C). ZTP reporter activity was observed in the efflux-deficient strain in response to 0.5 μg/mL TMP, increased in a dose-dependent manner until 2 μg/mL TMP, and remained stable at higher concentrations. In contrast, reporter activity was not detected for the isogenic efflux-proficient strain at concentrations of TMP below 15 μg/mL (Figure 2C). Similar levels of *lacZ* activity were measured for the efflux-proficient strain treated with 100 μg/mL TMP as for the efflux-deficient strain exposed to 2 μg/mL TMP. As TMP is reported to be a substrate for the MexAB-OprM, MexCD-OprJ and MexEF-OprN efflux systems (28, 51-53), active efflux of TMP likely prevents folate cycle disruption and riboswitch activation at lower antibiotic concentrations, leading to the observed ∼50-fold difference in reporter sensitivity between the two strain backgrounds.

### Development and validation of a high throughput screening protocol

We next adapted the riboswitch-based reporter assay for high throughput screening (HTS) applications. The tube-based β-galactosidase Miller assay (54) was transferred to a 384-well plate format in which *P. aeruginosa* bacterial cells were efficiently lysed using a mixture of the commercial detergent PopCulture® and chicken egg white lysozyme (55, 56). β–Galactosidase activity was assayed using the substrate ONPG, with OD_420_ measured every minute for 30 minutes and calculated as the enzymatic rate normalized by the number of cells present in each well (Vmax/OD_600_).

A key requirement of a high throughput screening (HTS) protocol is the ability to differentiate true active compounds (hits) from noise with confidence (57, 58). Ideally, a reporter should exhibit low basal activity under non-inducing conditions, and high activity upon induction of the system. These traits ensure a high dynamic range of induction and minimize false positive hits. Using TMP as a positive control, we optimized bacterial growth phase for the 384-well format, varying both the duration of subculture (2-4 hours) from overnight cultures and the time of exposure to TMP (1-3 hours) (Figure 3).

**Figure 3.**
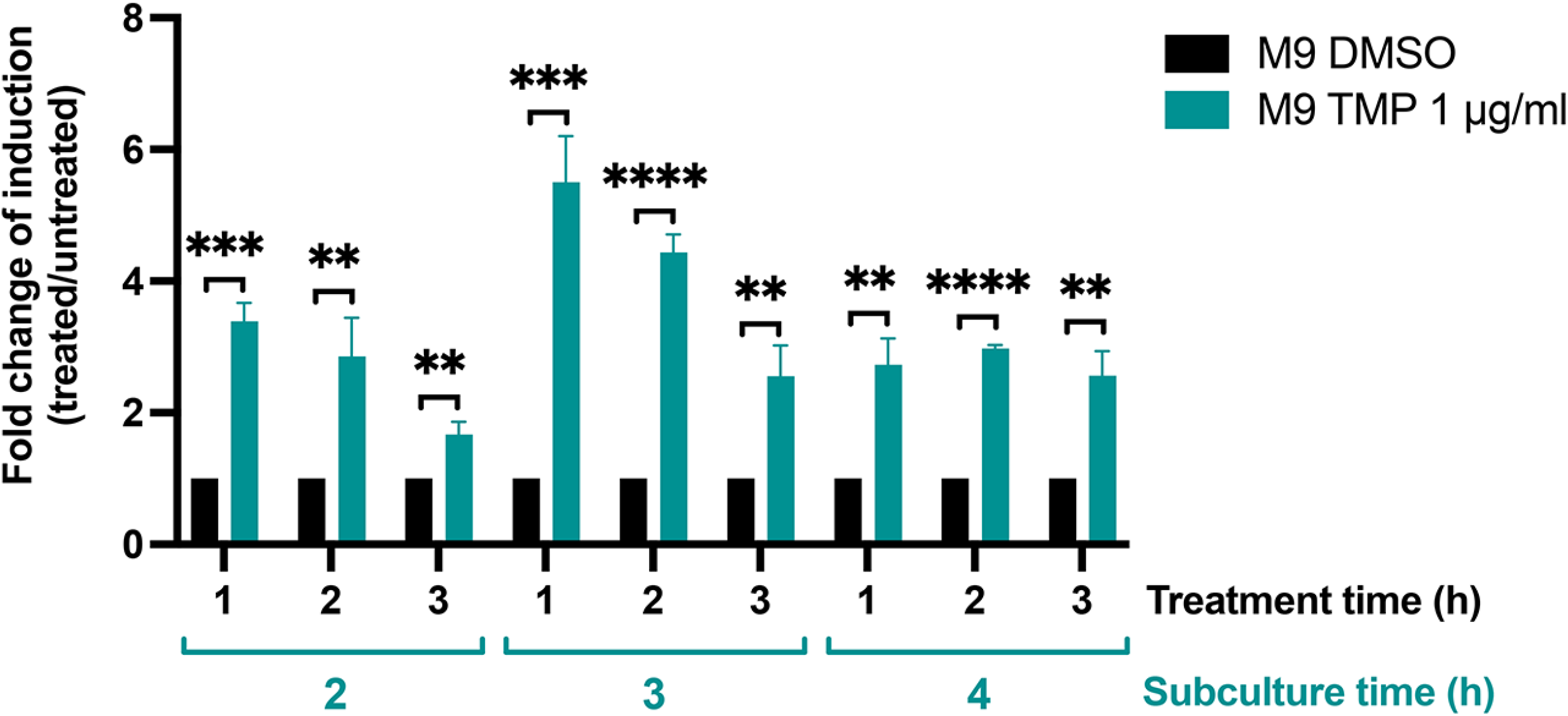
Adaptation of a β-galactosidase assay protocol to a 384-well plate HTS suitable format. The PAO1 Δefflux strain carrying the P*exoT-ZTP-lacZ* reporter was subcultured in M9 + 1% CAA for 2, 3 or 4 h in 125 mL flasks, then aliquoted into 384-well plates. Wells were treated with 1% DMSO (black histograms) or TMP 1 μg/mL (teal histograms). β-Galactosidase activity was measured after 1, 2 and 3 h of treatment and expressed as fold-change of induction values between cultures treated with TMP and cultures treated with the DMSO vehicle (1% final concentration) in each timepoint. Bars show mean ± standard deviation from 3 independent experiments. Unpaired two-tailed t-tests were used to compare β-galactosidase activity in cultures grown without vs. with TMP. P < 0.01, **; P < 0.001, ***; P < 0.0001, ****.

The highest fold-change of induction between cultures treated with TMP vs untreated controls was observed with 3 hours of subculture followed by 1 - 2 h of TMP exposure (Figure 3). Based on these results, we subcultured bacteria for 3h before exposing them for 1h to test compounds in subsequent experiments.

The Z’-factor screening coefficient described by Zhang and colleagues (58) assesses the ability of a HTS assay to identify true hits from a compound library. We determined the Z’-factor for our assay, using the 3 h subculture and 1 h treatment times established above, with TMP (1 μg/mL) and vehicle (1% DMSO) serving as positive and negative controls, respectively. Three independent trials, carried out in triplicate, yielded Z’-factor values of 0.605, 0.792, and 0.695, indicating that our HTS screening approach can reliably discriminate between a positive “hit” and background “noise” (0.5 ≤ Z’ < 1) (58).

### The ZTP riboswitch reporter is responsive to trimethoprim analogs

To further validate the functionality of the *PexoT*-ZTP-*lacZ* fusion in *P. aeruginosa*, we exposed our reporter strains to a small library of structurally diverse compounds with known mechanisms of action and distinct biological targets (Table 2). This library was enriched with dihydrofolate reductase and dihydropteroate synthetase inhibitors, as well as a range of other compounds currently in use in the clinic.

**Table 2.**
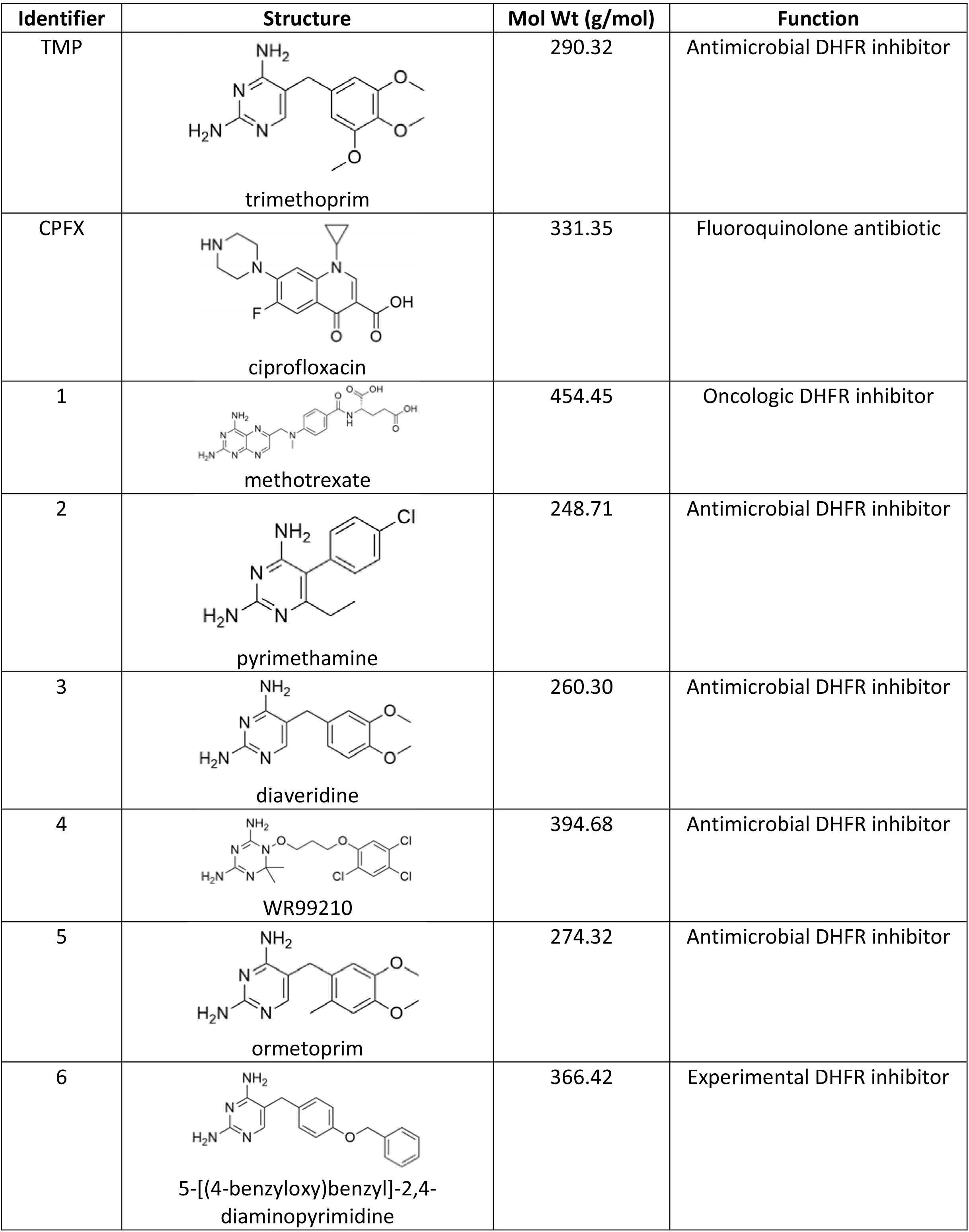
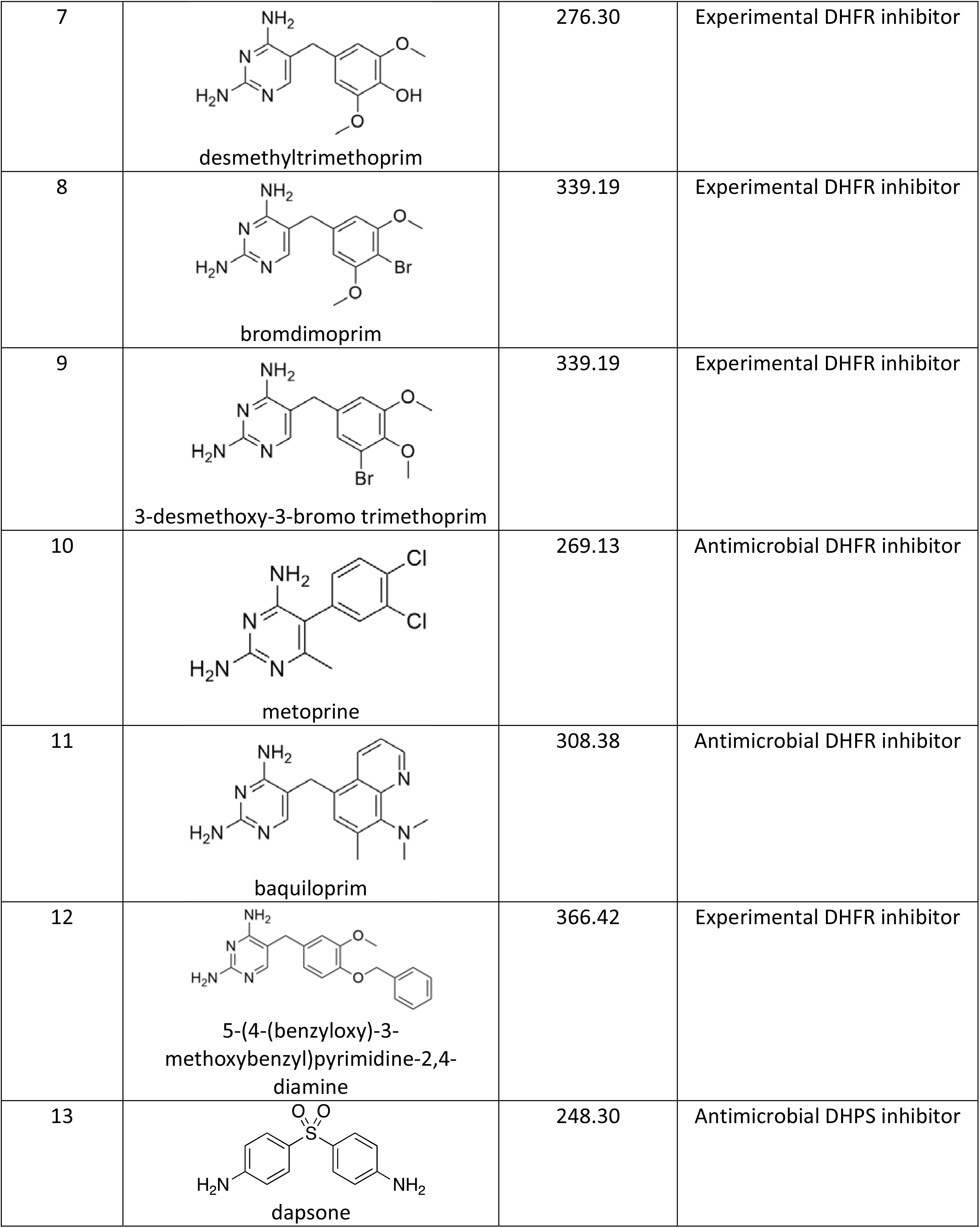
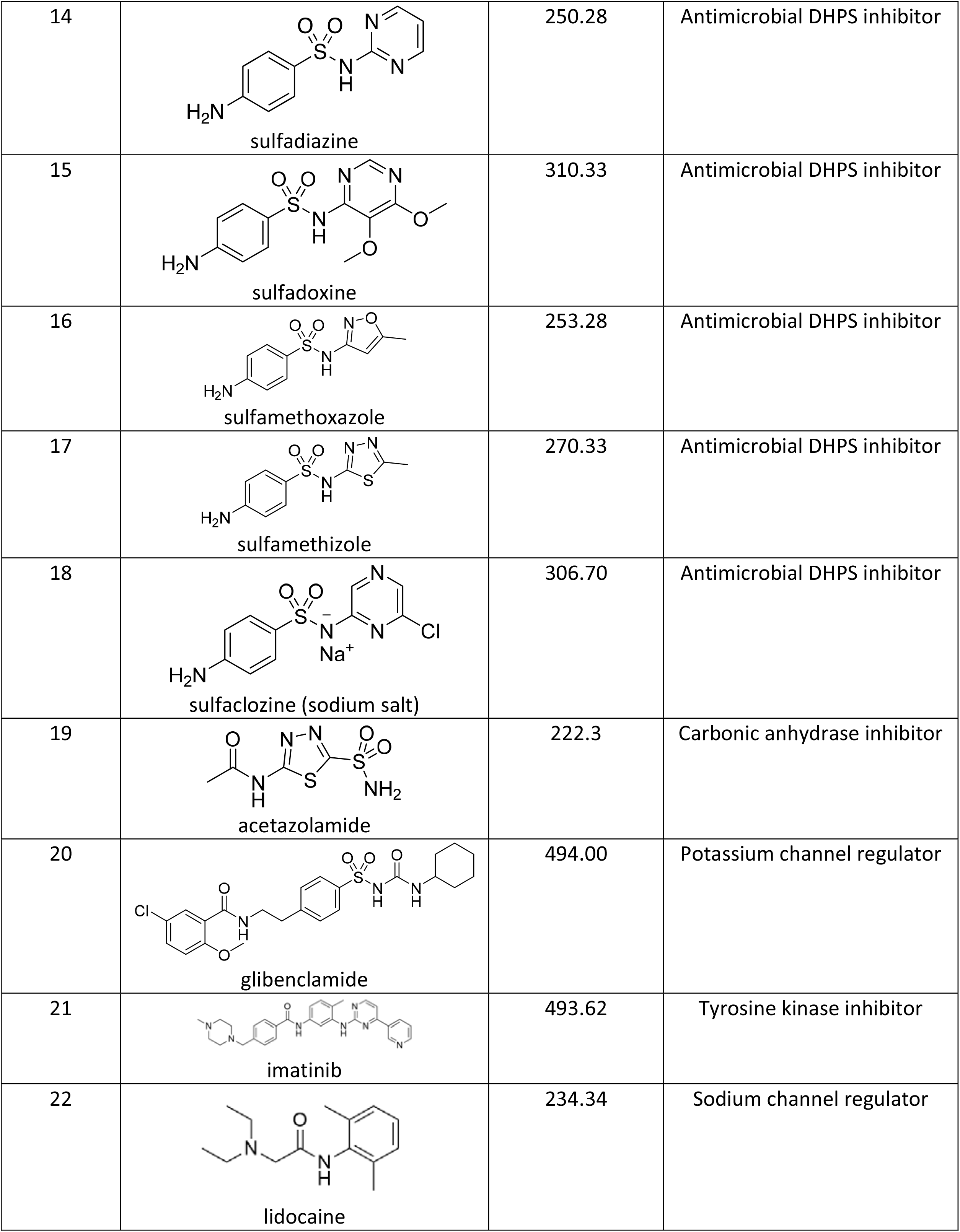
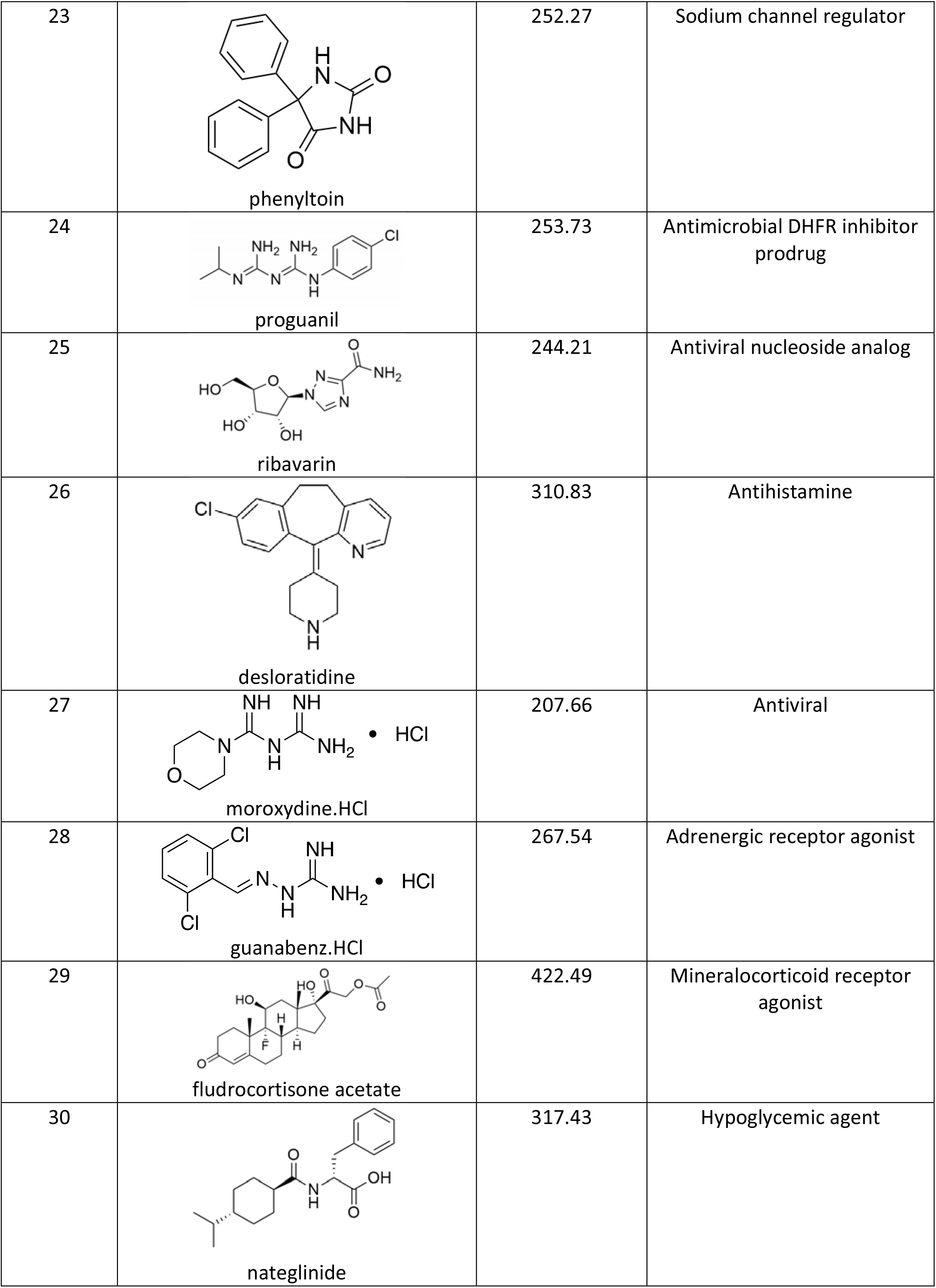
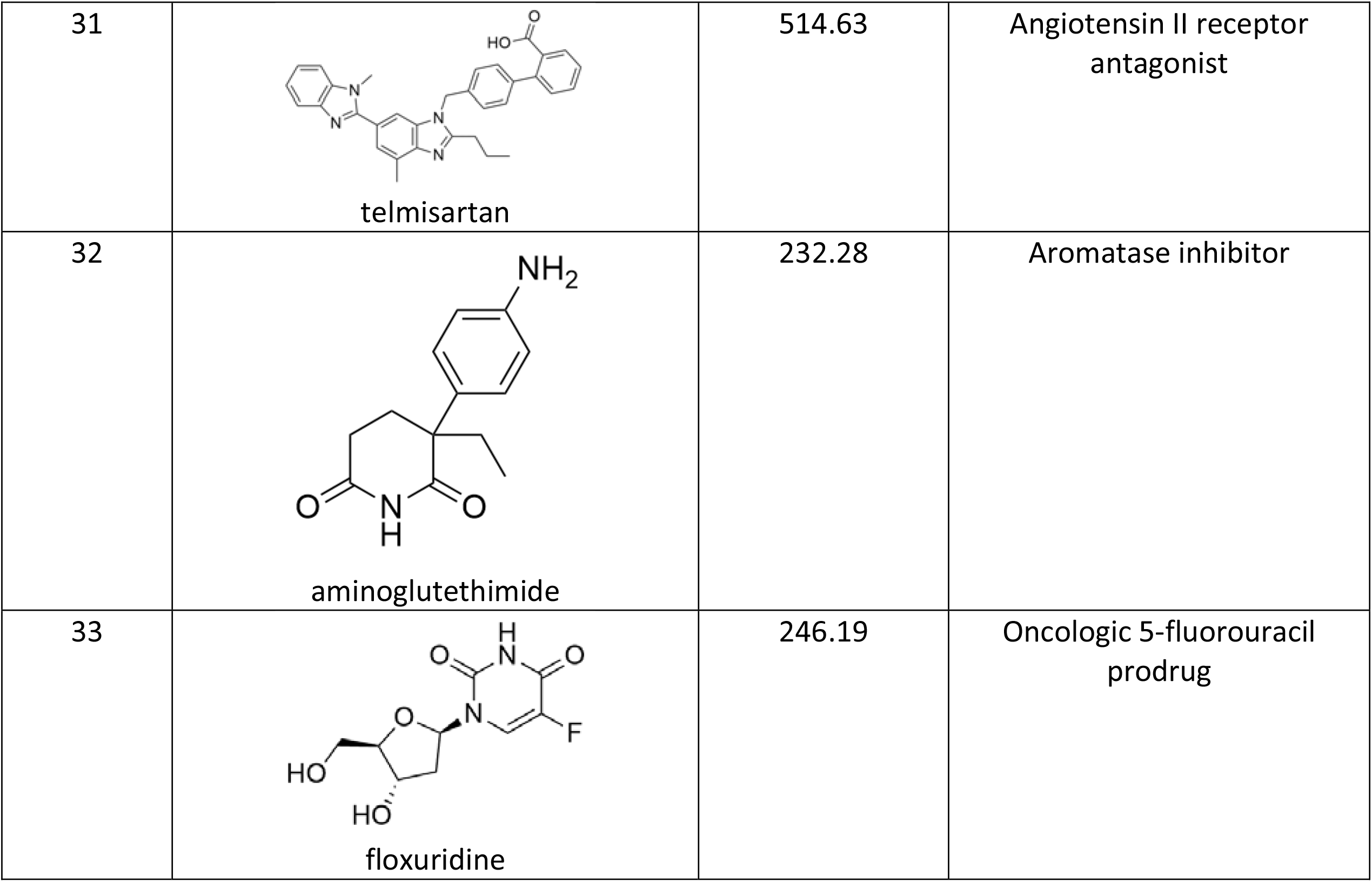
Structure, molecular weight, and function of the compounds tested in the ZTP-*lacZ* riboswitch reporter screen.

The PAO1 Δefflux strain carrying the *PexoT*-ZTP-*lacZ* reporter was treated with each compound at two concentrations, 10 and 50 μM. TMP was included as a positive control of induction (10 μM = 2.9 μg/mL and 50 μM = 14.5 μg/mL). Most tested compounds did not induce reporter expression and gave similar values of *lacZ* induction as the negative control (M9 + 1% CAA DMSO = 1, denoted as a dotted line in figure 4). Only typical dihydrofolate reductase inhibitors including the positive control TMP and compounds 2-5 and 7-12 induced reporter activity (Figure 4A). Notably, for TMP and WR99210 (Compound 4), reporter expression was higher at 10 μM than at 50 μM, suggesting that the higher concentration affected bacterial gene expression. Only methotrexate (Compound 1), known to have poor antibiotic activity against whole cells despite being a potent inhibitor of the isolated enzyme (59, 60), and compound 6 – with a disubstituted benzene ring substitution pattern that is atypical for DHFR inhibitors – were inactive among the diaminopyrimidines (Figure 4A). To confirm that reporter induction was exclusively due to ligand binding to the riboswitch aptamer, the positive hits identified in figure 4A were re-screened with the PAO1 Δefflux strain carrying the *PexoT*-ZTP(M4)-*lacZ* riboswitch reporter variant defective for ZTP binding. The absence of *lacZ* induction in this strain (Figure 4B) confirmed that these compounds act by disrupting the folate pathway in *P. aeruginosa* and increasing ZTP/ZMP concentrations. All nominal dihydropteroate synthetase inhibitors – including dapsone (Compound 13) and sulfamethoxazole (Compound 16) - were inactive, both at the screening concentrations and when subsequently tested at 250 μM and 500 μM (Figure 4C). Lastly, all compounds that are not known folate pathway inhibitors – acetazolamide, glibenclamide, imatinib, lidocaine, phenyltoin, proguanil, ribavirin, desloratadine, moroxydine, guanabenz, fludrocortisone acetate, nateglinide, telmisartan, aminoglutethimide, and floxuridine – were inactive in this assay.

**Figure 4.**
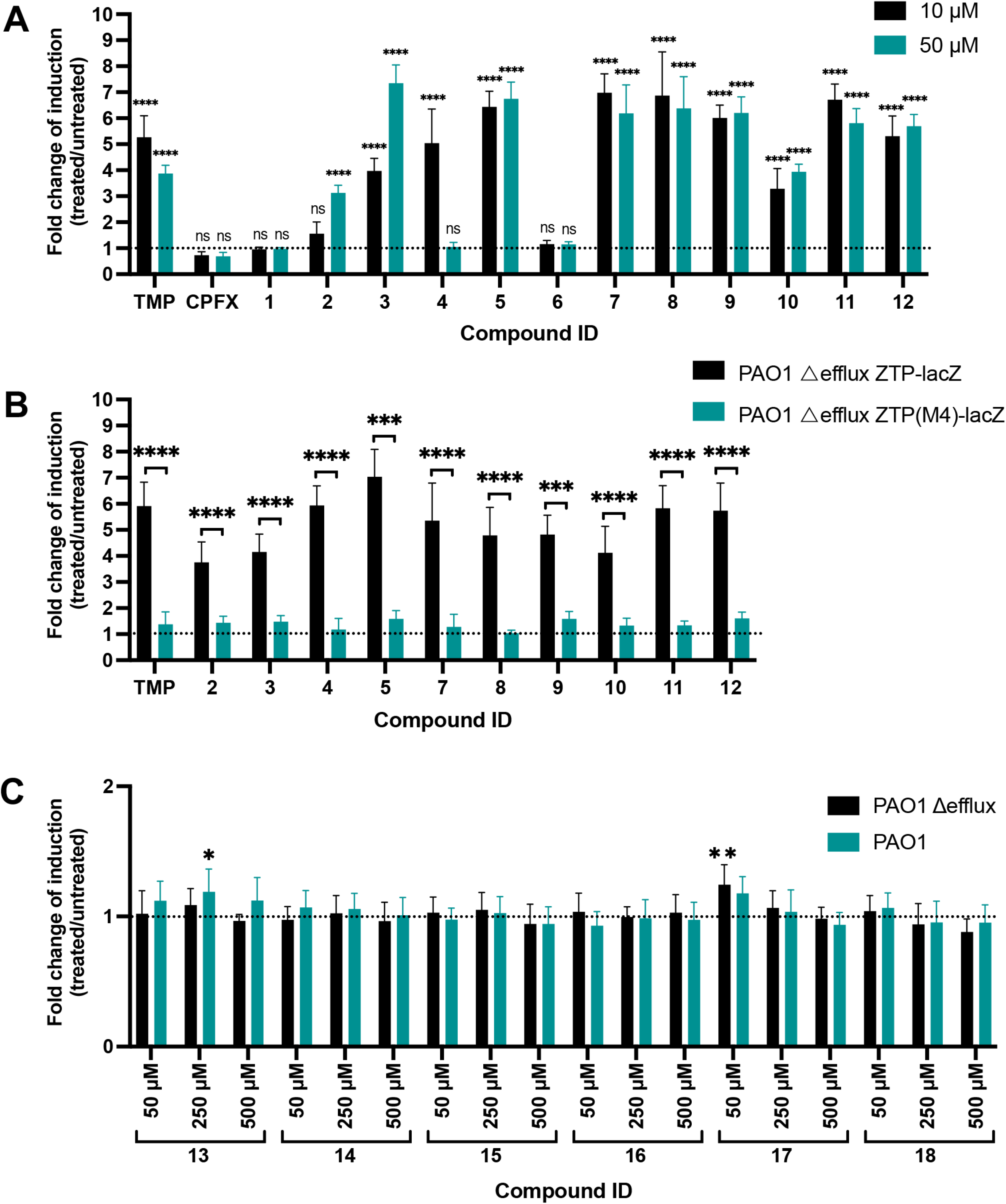
Response of the ZTP riboswitch reporter to bioactive molecules. **A**. β-Galactosidase activity of the PAO1 Δefflux Δ*exsD* P*exoT*-ZTP-*lacZ* exposed to a focused library of dihydrofolate reductase inhibitors at a final concentration of 10 and 50 μM (Table 2). The basal level of β-galactosidase activity of cultures grown in M9 + 1% CAA 1% DMSO (control) is represented by the dotted line. Values are averages and standard deviations from 3 independent experiments. One-way ANOVA coupled with Dunnett’s multiple comparison test were employed to compare the β-galactosidase activity values of cultures exposed to each compound (10 μM, black histograms; 50 μM, teal histograms) versus the control (M9 + 1% CAA 1% DMSO, dotted line). P < 0.0001, ****; not significant, ns. **B**. β-Galactosidase activity of PAO1 Δefflux Δ*exsD* P*exoT*-ZTP-*lacZ* (black histograms) and PAO1 Δefflux Δ*exsD* P*exoT*-ZTP(M4)-*lacZ* (teal histograms) exposed to the hits identified in 4A at a final concentration of 50 μM for most compounds except compounds 4 and TMP, which were assayed at 10 μM. The basal level of β-galactosidase activity of cultures grown in M9 + 1% CAA 1% DMSO (control) is represented by the dotted line. Values are averages and standard deviations from 3 independent experiments. Unpaired two-tailed t-tests were used to compare the β-galactosidase activity values of PAO1 Δefflux Δ*exsD* P*exoT*-ZTP-*lacZ* and PAO1 Δefflux Δ*exsD* P*exoT*-ZTP(M4)-*lacZ* cultures exposed to each of the compounds. P < 0.001, ***; P < 0.0001, ****. **C**. β-Galactosidase activity of the ZTP riboswitch reporter in the efflux-deficient (black histograms) and efflux-proficient (teal histograms) *P. aeruginosa* backgrounds to different concentrations of dihydropteroate synthetase inhibitor drugs present in the compound library (Table 2). The dotted line represents the basal level of β-galactosidase activity of cultures grown in M9 + 1% CAA DMSO (control). Values are averages and standard deviations from 3 independent experiments. One-way ANOVA coupled with Dunnett’s multiple comparison test were employed to compare the β-galactosidase activity values of cultures of efflux-deficient (black histograms) and efflux-proficient strains (teal histograms) exposed to different concentrations of each compound vs. the control (M9 + 1%CAA 1% DMSO, dotted line). P < 0.05, *; P < 0.01, **; absence of asterisks means that the difference was not significant.

### The ZTP riboswitch reporter system can distinguish compounds that are subject to efflux versus those that are retained inside cells

The ZTP riboswitch reporter, expressed in efflux-deficient bacteria, identified several dihydrofolate reductase inhibitors capable of crossing the bacterial cell envelope and disrupting purine synthesis. To determine if the activity of these compounds was affected by the presence of efflux systems, we repeated our screen using the efflux-proficient strain carrying the ZTP riboswitch (Figure 5A). In most instances, β-galactosidase expression was diminished in the efflux-proficient cells relative to their efflux-deficient counterparts, suggesting that these compounds were substrates for efflux. Three compounds, however, continued to show values of fold change of induction of β-galactosidase expression ≥ 3 in the efflux-proficient strain: WR99210 (Compound 4), desmethyltrimethoprim (Compound 7), and baquiloprim (Compound 11). We employed the *PexoT*-ZTP(M4)-*lacZ* mutant riboswitch construct to confirm that *lacZ* induction was dependent on ZTP binding to the riboswitch aptamer, allowing us to conclude that these compounds targeted the folate pathway in the efflux-proficient background (Figure 5B).

**Figure 5.**
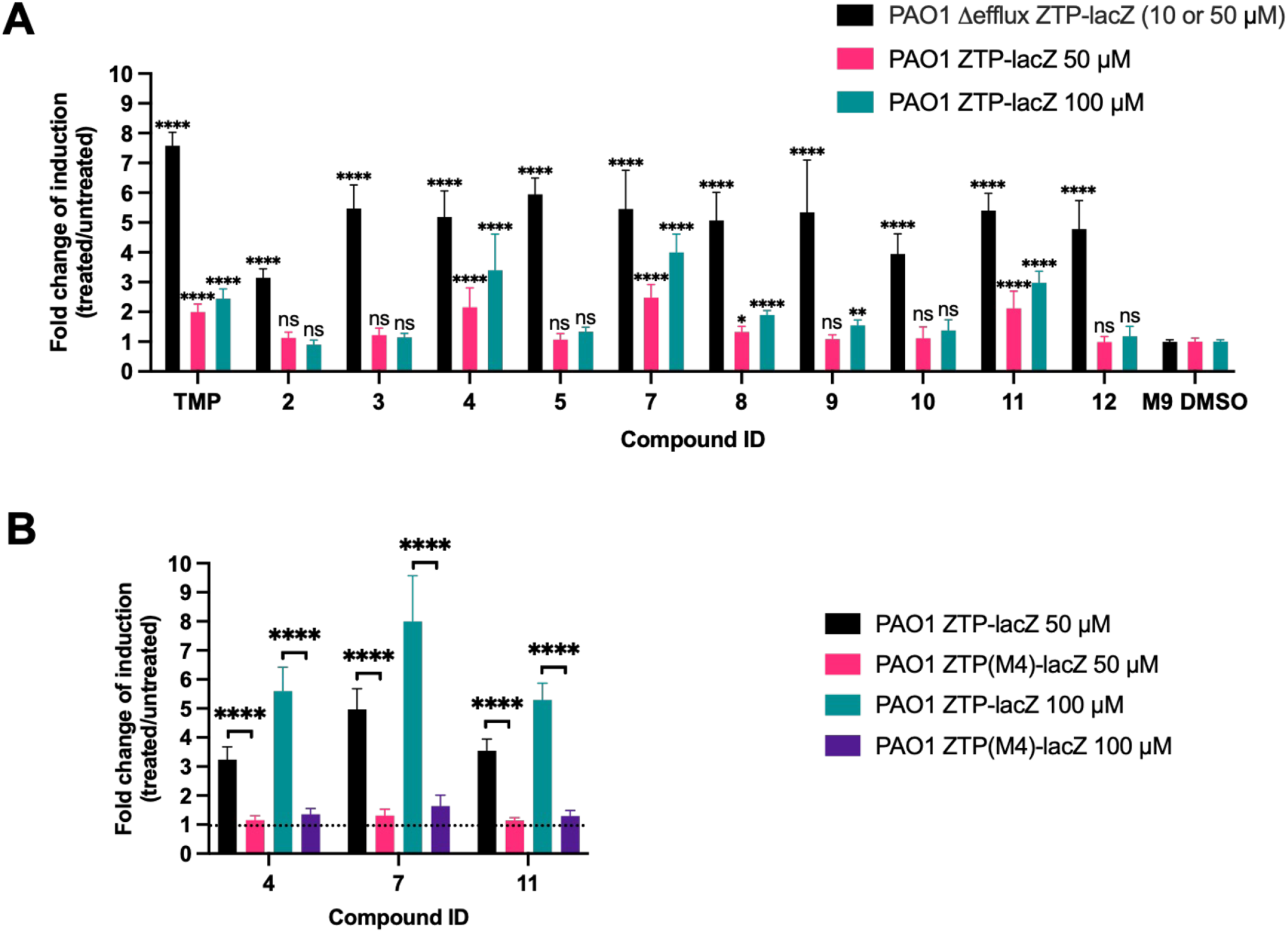
Response of the ZTP riboswitch reporter against dihydrofolate reductase inhibitors in the efflux-proficient background. **A**. Comparison of β-galactosidase activity levels of PAO1 Δefflux Δ*exsD* P*exoT*-ZTP-*lacZ* and PAO1 Δefflux Δ*exsD* P*exoT*-ZTP-*lacZ* exposed to a focused compound library (Table 2). For the efflux deficient strain (black histograms), most compounds were used at a final concentration of 50 μM, except for compounds 4 and TMP, which were used at a final concentration of 10 μM. The efflux-proficient strain (magenta and teal histograms) was treated with both 50 and 100 μM of each compound. Values are averages and standard deviations from 3 independent experiments. One-way ANOVA coupled with Dunnett’s multiple comparison test were employed to compare the β-galactosidase activity values of cultures treated with each of the compounds (black histograms for the efflux-deficient strain; magenta and teal histograms for the efflux-proficient strain) with the control (M9 + 1% CAA 1% DMSO, labelled M9 DMSO). P < 0.05, *; P < 0.01, **; P < 0.0001, ****; not significant, ns. **B**. β-Galactosidase activity of PAO1 Δ*exsD* P*exoT*-ZTP-*lacZ* (black and teal histograms), and PAO1 Δ*exsD* P*exoT*-ZTP(M4)-*lacZ* (magenta and violet histograms) exposed to the hits identified in 5A at final concentrations of 50 μM and 100 μM. The dotted line represents the basal level of β-galactosidase activity of cultures grown in M9 + 1% CAA 1% DMSO (control). Values are averages and standard deviations from 3 independent experiments. Unpaired two-tailed t-tests were used to compare the β-galactosidase values of PAO1 Δ*exsD* P*exoT*-ZTP-*lacZ* with PAO1 Δ*exsD* P*exoT*-ZTP(M4)-*lacZ* cultures exposed to the compounds from the library in final concentrations of 50 or 100 μM. P < 0.0001, ****.

We hypothesized that the measured differences in *lacZ* induction between efflux-deficient vs. efflux-proficient bacteria reflected the relative retention of folate cycle-disrupting compounds in these otherwise isogenic bacterial strains. To test this directly, we extracted the cell-internalized compounds from efflux-proficient and efflux-deficient *P. aeruginosa* strains treated with 50 μM of either ormetoprim (Compound 5) or desmethyltrimethoprim (Compound 7) and measured their relative abundance by high-resolution liquid chromatography-mass spectrometry (LC-MS) (Figure 6). The internal abundance of ormetoprim was greater in the efflux-deficient strain relative to the efflux-proficient strain, in good agreement with the ability of this compound to induce *lacZ* expression solely in the efflux-deficient reporter strain. In contrast, the internal abundance of desmethyltrimethoprim – which induces *lacZ* expression to a similar extent in both efflux-proficient and efflux-deficient reporter strains – did not differ significantly between these two backgrounds. These results suggest that the ability of a compound to induce *lacZ* expression in the ZTP riboswitch reporter system reflects both its ability to cross the cell envelope and to avoid active efflux.

**Figure 6.**
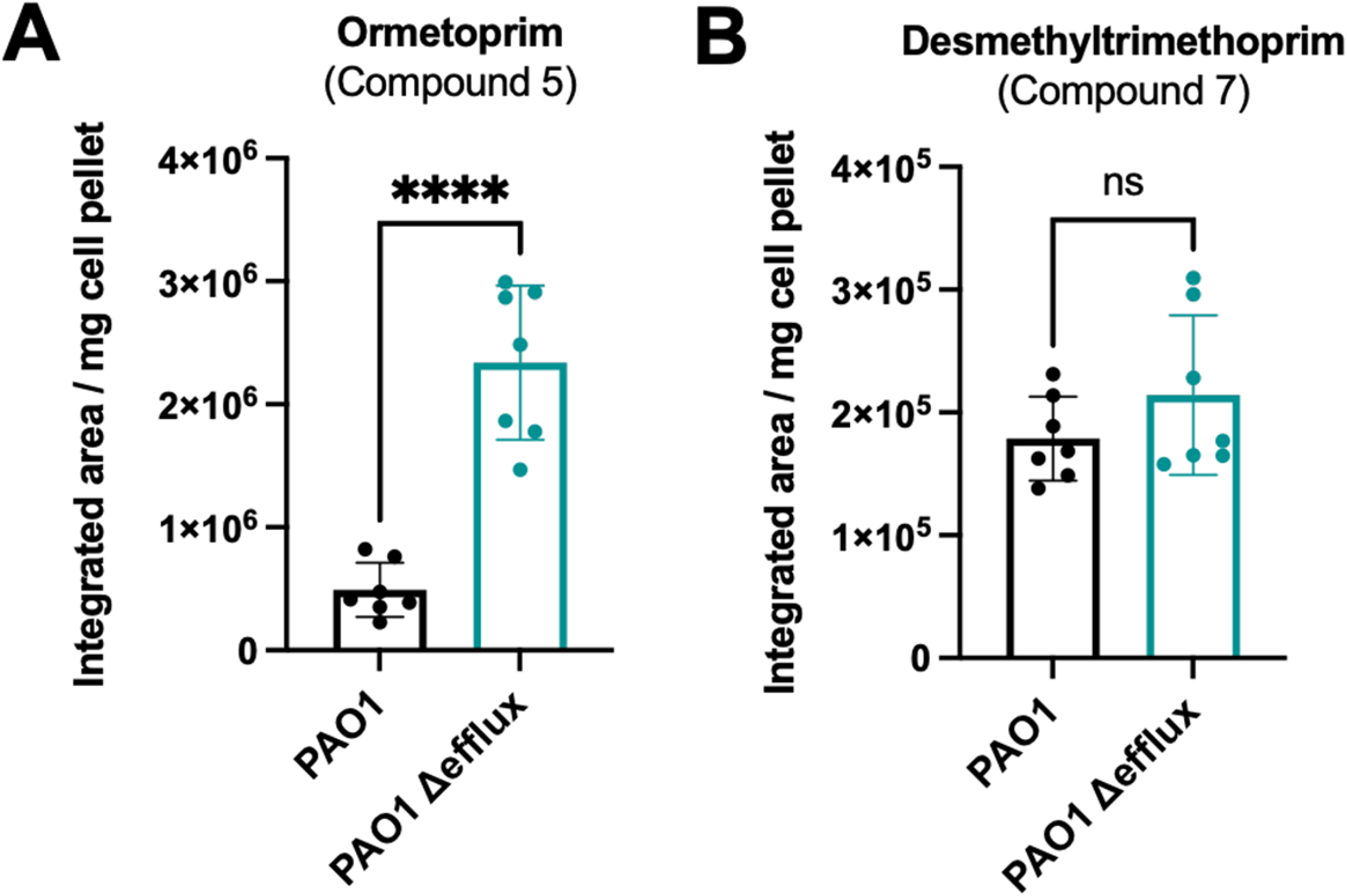
Relative internal abundances of select compounds in efflux-deficient and efflux-proficient *P. aeruginosa* strains normalized to dry cell pellet weight. Patterns of compound accumulation in efflux-proficient (black histograms) and efflux-deficient (teal histograms) strains treated with an actively exported compound (A, ormetoprim), and a compound that enters the cells and is not exported (B, desmethyltrimethoprim). Histograms showing the mean values of 7 biological replicates along with standard deviation error bars are plotted. Unpaired two-tailed t-tests were used to compare the normalized values. P < 0.0001, ****; not significant, ns.

## Discussion

We present an experimental approach to identify folate pathway inhibitors that cross the bacterial envelope of *P. aeruginosa* and to classify them as substrates for efflux systems. This approach is robust enough to undertake a high throughput screening campaign to elucidate the dynamics of compound uptake, retention, and efflux in this Gram-negative bacterium. The screening system employs a previously reported ZTP riboswitch-based reporter (32, 38) adapted to *P. aeruginosa*, with the reporter inserted in single-copy at a silent chromosomal location (*attB)*(61) and expressed constitutively and uniformly from a type three secretion system promoter (43) (Figure 2A). The strength of our approach lies in the use of this reporter system in two isogenic backgrounds: an efflux-proficient PAO1 wild-type strain and an isogenic efflux-deficient mutant lacking multiple efflux systems, i.e. MexAB-OprM, MexCD-OprJ, MexJK, MexXY and MexEF-oprN (48). These efflux systems belong to the resistance nodulation division (RND) family and are important contributors to the efflux of noxious compounds (28, 62-64). In the efflux-deficient strain, the reporter is activated by the control drug trimethoprim (TMP) at a concentration ca. 50 times less than that required to observe a similar effect in the efflux-proficient strain (Figure 2C). These results are consistent with previous reports that identified TMP as substrate of the MexAB-OprM, MexCD-OprJ and MexEF-OprN efflux systems in *P. aeruginosa* (reviewed in (8, 28, 65)). Most importantly, this observation suggests that this reporter system can (i) identify compounds that cross the bacterial envelope, and (ii) distinguish between compounds that remain inside the cell vs those that are efflux substrates.

We subsequently adapted this reporter system for high throughput screening (HTS) of compounds in 384-well plate format. The method requires few hands-on manipulations, is straightforward to automate, and allows thousands of compounds to be screened per day. Although HTS assays employing *lacZ*-based reporters often rely on fluorogenic substrates such as 4-methylumbelliferyl-β-D-galactopyranoside (4-MUG) (31, 32), this molecule does not cross the envelope of *P. aeruginosa* and requires cell lysis, while the high intrinsic fluorescence of *P. aeruginosa* PAO1 markedly reduces the S/N ratio of 4-MUG. We devoted substantial effort to developing an efficient *P. aeruginosa* lysis method compatible with polystyrene 384-well plates and requiring minimal handling. Of the different approaches tested, which included freeze/thawing cycles and lysozyme treatment (66), we found treatment with the proprietary detergent-based PopCulture® reagent coupled with lysozyme-mediated lysis to be most efficient and most reproducible (Figure 3). The assay performed well with a modest period of outgrowth (3 hours of subculture) and gave a good response with only 1 hour of compound treatment (Figure 3). Under these conditions, the Z’-factor (58) ranged between 0.60 and 0.79, indicating a clear separation between the signals from the negative and positive controls (58).

A restrictive bacterial envelope and the action of numerous efflux systems are key determinants for *P. aeruginosa*’s increased basal resistance to many antimicrobials (8, 26-29, 67). One advantage of our screening approach is the utilization of a *P. aeruginosa* strain that lacks the main efflux systems responsible for antimicrobial export. As a consequence of these genetic modifications, this strain exhibits levels of antibiotic resistance that are more similar to the ones reported for *E. coli*. For instance, the PAO1 Δefflux strain exhibits a MIC of TMP of 15 μg/mL, a value closer to the reported 0.12-0.5 μg/mL of *E. coli* (68, 69) than to the 450 μg/mL observed for PAO1. The decrease in MIC is caused by the lack of efflux systems that can rapidly export TMP before it targets the folate pathway and inhibits bacterial growth. By integrating the ZTP riboswitch-based reporter into this efflux-deficient strain of *P*. aeruginosa, we could detect folate pathway disruption at much lower concentrations of TMP (Figure 2C), and potentially identify compounds that would appear “inactive” against an efflux-proficient strain. Further, the sensitivity of the riboswitch to perturbations of folate cycle homeostasis allowed compounds to be detected at concentrations far below those that affected viability – in the case of TMP, ∼30-fold below the MIC in both wild-type and efflux-deficient backgrounds (Figure 2C). This provides a significant increase in sensitivity over traditional bacterial viability assays, and the ability to identify permeating compounds even at the fixed (and relatively low) concentrations employed in high-throughput screens.

By employing a small library of bioactive small molecules enriched in folate pathway inhibitors we could determine if - in *P. aeruginosa -* the ZTP riboswitch reporter is responsive to other antifolate drugs. Dihydropteroate synthetase (DHPS) inhibitors, predominantly sulfonamides, impair the incorporation of *p*-aminobenzoic acid into dihydropteoric acid, and dihydrofolate reductase (DHFR) inhibitors, predominantly diaminopyrimidines, impair the reduction of dihydrofolic acid into tetrahydrofolic acid, both compounds targeting key steps in the biosynthesis of tetrahydrofolic acid (70, 71). Although the ZTP reporter in *E. coli* responds to both DHPS and DHFR inhibitors (32), the reporter in *P. aeruginosa* only showed increased expression when exposed to DHFR inhibitors (Figure 4A). No induction was observed when P. *aeruginosa* was exposed to the DHPS inhibitors in the library, which included most class members currently in use in the clinic. Efflux-mediated intrinsic resistance of *P. aeruginosa* to both sulfonamide and diaminopyrimidines drugs has been previously reported (52). In contrast with this previous report, our data suggest that in the case of sulfonamides, such resistance is mostly mediated by poor permeation rather than by the action of active efflux, as efflux-deficient and -proficient strains showed similar expression levels of the reporter when challenged with sulfonamides, even at high concentrations (Figure 4C).

Most compounds that strongly induced the ZTP riboswitch reporter in the efflux-deficient strain had minimal effect on the efflux-proficient strain (Figure 5A), suggesting that these molecules were substrates for active efflux. This included DHFR inhibitors which see concurrent clinical use – albeit typically not for *P. aeruginosa* infection – such as trimethoprim (TMP), pyrimethamine (Compound 2), diaveridine (Compound 3), ormetoprim (Compound 5), and metoprine (Compound 10). In contrast, three compounds in our library appeared to be exported less efficiently than the others: WR99210 (Compound 4), desmethyltrimethoprim (Compound 7), and baquiloprim (Compound 11) (Figure 7). This riboswitch-based observation was validated by LC-MS experiments that measured the relative intracellular abundance of a compound that our riboswitch assay categorized as an efflux substrate – ormetoprim (Compound 5) - or non-efflux substrate – desmethyltrimethoprim (Compound 7). While it remains possible that WR99210, desmethyltrimethoprim or baquiloprim are simply poor up-regulators of efflux, we think this less likely given the brief period of compound exposure (1h) in our assay. DHFR inhibitors have not been reported to induce efflux system expression in *P. aeruginosa* and only two efflux pumps have been identified to be induced by antimicrobial agents: MexXY by ribosome-binding inhibitors like tetracycline and chloramphenicol (72, 73), and MexCD-OprJ by membrane damaging agents like chlorhexidine and polymyxin B (74).

**Figure 7.**
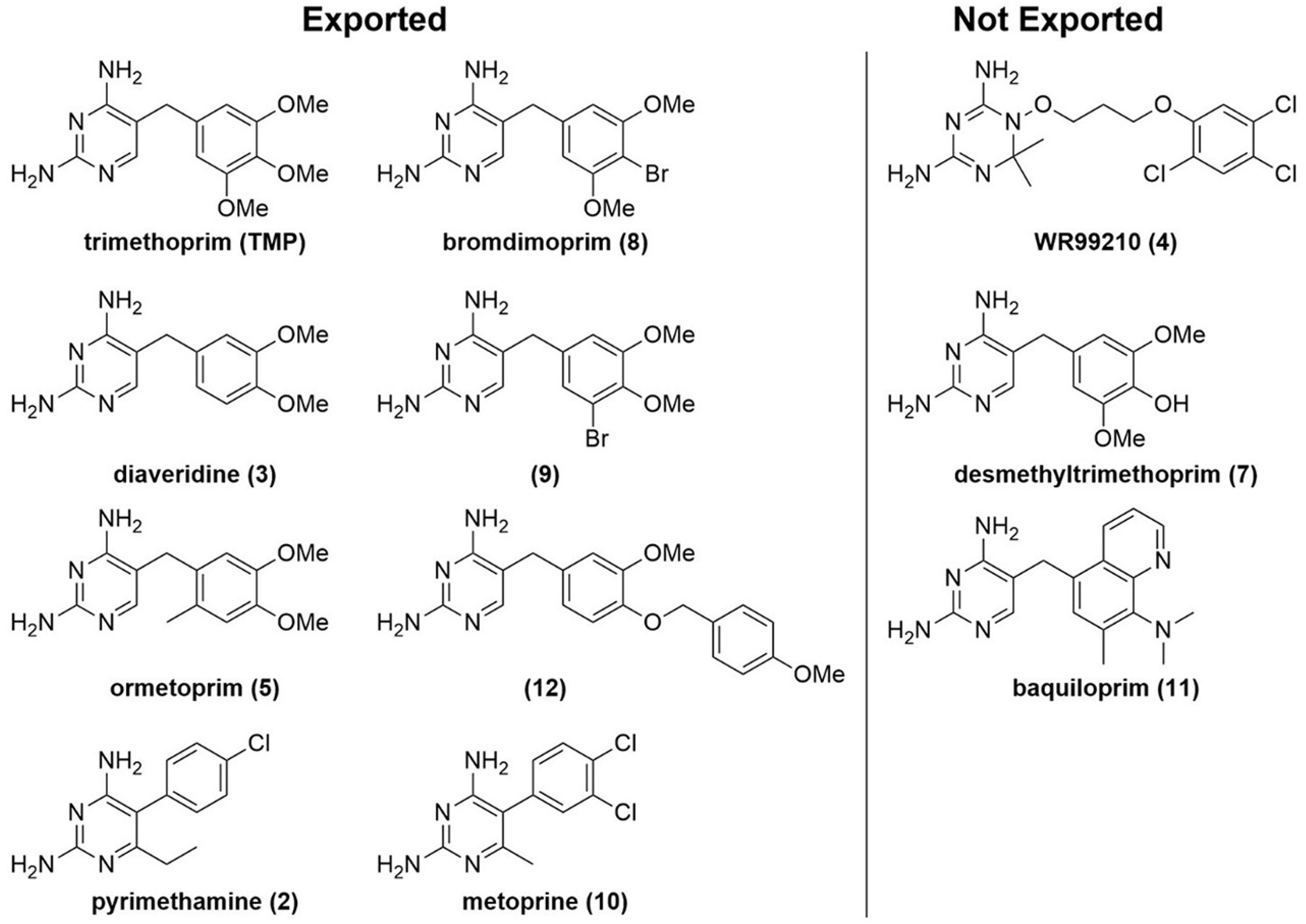
DHFR inhibitors that enter *P. aeruginosa* classified by exported through active efflux (left) and remaining inside the cell (right).

Consideration of the physical properties of the compounds that are not exported reveals no obvious trends relative to those that are. WR99210, initially developed as a potential antimalarial (75) but seeing no clinical use, has a calculated LogP (cLogP) of 2.47 and total polar surface area (TPSA) (95 Å^2^) well within the ranges of those exported (cLogP 0.98-3.07, TPSA 77-104 Å^2^). Similarly, baquiloprim, used as a veterinary anti-infective (76), has a cLogP of 2.64 and TPSA of 92 Å^2^. While desmethyltrimethoprim, recently noted as impeding antibiotic resistance evolution (77), is more polar (cLogP 0.65, TPSA 115 Å^2^) the difference from trimethoprim (cLogP 0.98, TPSA 104 Å^2^) is relatively small. It is therefore likely that differentiation between exported and not-exported compounds occurs at the protein-ligand interaction level.

We predict that screening thousands of compounds with this approach will lead to the discovery of numerous permeable compounds that are minimally exported by efflux systems in this Gram-negative bacterium, thus allowing investigators to identify pharmacophores and physicochemical properties that determine if a given compound will be exported by *P. aeruginosa*’s most important efflux systems or if it will remain inside the cell. Identification of such structural properties can inform the design of drugs able to bypass two of the most efficient defenses against antimicrobial compounds in Gram-negative bacteria: the restrictive nature of the bacterial envelope and the action of efflux pumps (Reviewed in (8, 9, 11, 12, 14, 63, 78, 79)).

## Materials and Methods

### Bacterial strains, media, culture conditions and chemicals

Bacterial strains and plasmids used in this study are listed in Table 1. Lysogeny broth (LB) was routinely used as liquid medium for both *E. coli* and *P. aeruginosa*. M9 minimal medium supplemented with 1% Casamino acids (M9 + 1% CAA) was used in specific cases. LB plates contained agar at 15 g/L final concentration. Vogel-Bonner minimal medium (80) plates (15 g/L agar) were used to select *P. aeruginosa* in specific cases. Cultures were grown at 37°C with shaking at 250 rpm in a New Brunswick Innova 44/44R shaker. When required, antibiotics were added at the following concentrations: for *E. coli*, 15 μg/mL gentamicin, 10 μg/mL tetracycline; for *P. aeruginosa*, gentamicin (30 μg/L for the efflux-deficient strain or 100 μg/mL for the efflux-proficient strain), tetracycline (20 μg/mL for the efflux-deficient strain or 200 μg/mL for the efflux-proficient strain), and carbenicillin (75 μg/mL for the efflux-deficient strain or 200 μg/mL for the efflux-proficient strain). All bacterial strains were stored at -80°C as 15% (v/v) glycerol stocks. Trimethoprim (TMP), used as a positive control for the ZTP riboswitch reporter, was purchased from MP biochemicals and dissolved in DMSO. The complete library of compounds selected for this study was purchased from Cayman Chemical, MedChemExpress, Ambeed, Toronto Research Chemicals, Ambinter, and Sigma-Aldrich. They are listed in Table 2 with molecular weight and function; these compounds were also dissolved in DMSO. 20% Sodium dodecyl sulphate (SDS) solution was purchased from AmericanBIO. The chromogenic substrate ortho-nitrophenyl β-D-galactopyranoside (ONPG) and chicken egg white lysozyme were purchased from Sigma-Aldrich. PopCulture® Reagent was purchased from Millipore.

### Strain construction

PCR primers employed for strain construction and verification were synthesized by the Keck facility (Yale University) and are listed in Table 1. Unmarked, in-frame deletion of ***exsD*** was carried out in PAO1 and its isogenic strain PA0397 (48) via allelic exchange as described in (81). Briefly, primers were designed to target the genomic regions upstream (Up-F and Up-R) and downstream (Down-F and Down-R) of the desired deletion (see Table 1). *attB1* and *attB2* sites were added to the 5’ ends of the Up-F and Down-R oligos, respectively. Up-R and Down-F oligos were designed to be complement of one another. Phusion polymerase (NEB) was used to amplify the regions flanking the *exsD* gene, which were then spliced together by overlap extension PCR (SOE-PCR), generating a linear DNA fragment containing the desired deletion. This DNA fragment was cloned into the Gateway suicide vector pDONRX (82) and verified by sequencing using primers attL1-F and attL2-R (Table 1). The resulting plasmid was transformed into *E. coli* S17-1 and mobilized into *P. aeruginosa* by mating. *Pseudomonas aeruginosa* merodiploids were selected on VBM plates with either 30 or 100 μg/mL gentamicin, depending on whether the recipient strain was efflux-deficient (PA0397) or efflux-proficient (WT). A second recombination event, loss of vector backbone, was selected by streaking merodiploids on VBM plus sucrose (10%). Δ*exsD* candidates that were both gentamicin and sucrose sensitive were isolated and screened by PCR using primers external to the deleted region (denoted E1 and E2, see Table 1) and confirmed by Sanger sequencing.

A DNA fragment consisting of the 112 bp promoter region of *exoT* (44), the 83 bp ZTP riboswitch sequence from *Pectobacterium carovotorum subsp. Carovotorum PC1* plus the first 17 nt of the *rhtB* gene coding sequence (38), and the complete *lacZ* gene (3078 bp) was synthesized (Genewiz Inc.) and cloned into mini-CTX2, allowing for integration into the *attB* site of the *P. aeruginosa* chromosome (61). This mCTX2-P*exoT-*ZTP-*lacZ* plasmid was transformed into *E. coli* S17-1 and mobilized by mating into *P. aeruginosa*. Integrants were selected on VBM plates with 20 or 200 μg/mL tetracycline, depending on whether the recipient strain was efflux-deficient (PA0397) or efflux-proficient (PAO1 or PA14 WT). The mCTX2 vector backbone was excised by mating with *E. coli* SM10 carrying the pFLP2 vector, which expresses Flp recombinase (83). Successful loss of vector backbone and pFLP2 was confirmed by PCR using *attB*-SER-F and neolacZ primers (Table 1) for candidates that were both tetracycline and carbenicillin sensitive.

A previously described mutated version of the ZTP riboswitch (M4) (38) was constructed by PCR amplification of the mCTX2-P*exoT-*ZTP-*lacZ* plasmid using the three GA primer pairs (Table 1) followed by NEBuilder assembly (NEB). The resulting P*exoT-*ZTP(M4)-*lacZ* reporter fusion was integrated in the *P. aeruginosa* chromosome as described above and confirmed by PCR and Sanger sequencing.

### β-galactosidase assays

Overnight cultures of *P. aeruginosa* carrying the riboswitch reporter fusion were diluted 1:100 into 2.5 mL of M9 + 1% CAA medium and incubated in 14 mL tubes with aeration at 37°C until early exponential phase (2 h, OD_600_ ≈ 0.06-0.1). At this time, cultures were treated with TMP over a range of concentrations (0.5 to 300 μg/mL). The final concentration of DMSO was 1%. Colorimetric β-galactosidase assays were performed as previously described (84). Briefly, two 200 μL aliquots of each culture were collected: one was employed to determine OD_600_, while the second aliquot was permeabilized in 1.5 mL microcentrifuge tubes containing 600 μL of Z-Buffer (60 mM Na_2_HPO_4_.7H_2_O, 40 mM NaH_2_PO_4_.H_2_O, 10mM KCl, 1mM MgSO_4_.7H_2_O, pH 7), 15 μL of 0.01% SDS and 30 μl Chloroform. The β-galactosidase enzymatic reaction was initiated by adding 200 μL ONPG 4 mg/mL as substrate; to stop the reaction, 500 μL of Na_2_CO_3_ 1M was added. 200 μL of the reaction supernatant was taken to measure OD_420_ and OD_550_. Enzymatic activity was determined using the following formula: LacZ activity = [(OD_420_ x 1000) – (1.75 x OD_550_)]/[OD_600_ x Reaction time (min) x Culture volume (mL)] (84).

### Determination of minimal inhibitory concentrations (MIC) of trimethoprim

*P. aeruginosa* was grown with aeration in M9 + 1% CAA to mid-log, then samples containing ∼ 1 × 10^5^ colony-forming-units (CFUs) were transferred to 14 mL tubes containing known concentrations of trimethoprim. After 16 h of incubation with aeration at 37°C, growth was scored (68).

### 384-well plate-based β-galactosidase assay (development of a high throughput screening protocol)

Overnight cultures of the PAO1 Δ*exsD* Δefflux *attB::*P*exoT*-ZTP-*lacZ* strain were diluted 1:100 in 20 mL of fresh M9 medium +1% CAA and incubated in 125 mL flasks with aeration at 37°C for 2, 3 or 4h (OD_600_ ≈ 0.06, ≈ 0.2, or ≈ 0.6, respectively). At this point, the cultures were divided in two. One fraction was treated with 1 μg/mL (or 3.4 μM) TMP dissolved in DMSO. The vehicle DMSO was added to the second fraction (negative control). 30 μL of each of the two fractions were aliquoted in 384-well plates and incubated at 37°C with aeration for 1, 2 or 3h. After each exposure time, OD_600_ was recorded using a Tecan Infinite 200 Pro plate reader. Subsequently, bacteria were lysed by adding a mixture of 4.5 μl of PopCulture® reagent and 0.5 μl of lysozyme 4 U/μl in each well, the mixture was incubated at room temperature for 20 min with 750 rpm shaking to guarantee a more efficient lysis. Following bacterial lysis, 40 μl of 1.46 mg/mL ONPG substrate dissolved in Z-Buffer were added to each well. Immediately after, the plate was placed in the plate reader prewarmed at 28°C, and OD_420_ reads were taken each minute for 30 min. A 5 s shaking step (1 mm amplitude linear shaking) was set between readings.

LacZ activity was determined following a previously described protocol (66), with some modifications. Briefly, the increase of OD_420_ in time (minutes) was plotted for each well, and the reaction rate (Vmax) was determined by extrapolation of the slope from the most linear part of the curve. These values were then normalized to the OD_600_ recorded for each well before bacterial lysis. LacZ activity values were expressed in Vmax/OD_600_ units. To estimate the values of fold-change of induction, the absolute LacZ activity values from the wells treated with TMP were divided by the values from the wells treated with DMSO. Z’-factor calculation was done as described by Zhang and colleagues (58).

### Screening of a focused library of compounds

To screen the focused collection of compounds, overnight cultures of *P. aeruginosa* carrying the riboswitch reporter fusion were diluted 1:100 in 20 mL fresh M9 +1% CAA medium and incubated with aeration at 37°C for 3 h. At this point, 30 μl of the cultures was aliquoted into 384-well plates containing aliquots of each compound to a final concentration of 10 and 50 μM (for efflux-deficient strains), and 50 and 100 μM (for efflux-proficient strains). Subsequently, the plates were incubated at 37°C with aeration for 1 h. After incubation with each of the compounds, wells were treated as described above, with the exception that the efflux-proficient strain was lysed adding 9 μl PopCulture® and 1 μl Lysozyme 4 U/μl in each well (that is, twice the volume employed for the efflux-deficient strain), and lysis incubation time was 60 min. β-Galactosidase activity quantification of each well was calculated as before.

### Determination of intracellular levels of ormetoprim and desmethyltrimethoprim

Overnight cultures of the PAO1 Δ*exsD* Δefflux *attB::*P*exoT*-ZTP-*lacZ* and the PAO1 Δ*exsD attB::*P*exoT*-ZTP-*lacZ* strains were diluted 1:100 in 60 mL of fresh M9 medium +1% CAA and incubated in 250 mL flasks with aeration at 37°C until early exponential phase (3h, OD_600_ ≈ 0.2). At this point, the cultures were divided in three 20 mL fractions. These three fractions were treated with 50 μM solution of ormetoprim, 50 μM solution of desmethyltrimethoprim, or vehicle (1% DMSO), respectively. After 1 h incubation at 37°C with aeration, the cultures were pelleted and washed twice with 20 mL 1X PBS. After the second wash, the pellets were dried *in vacuo*, resuspended in 1 mL of a acetonitrile:methanol:water (2:2:1) solution and sonicated for 10 mins. The mixtures were centrifuged, and the supernatants transferred to glass vials and dried *in vacuo*. The dried samples were resuspended in 200 μl of a water:methanol (1:1) solution and 3 μl of each sample were subjected to high-resolution liquid chromatography-mass spectrometry (LC-MS) analysis (Agilent iFunnel 6550 quadrupole time-of-flight (QTOF), positive-mode electrospray ionization). Chromatography was performed on a Kinetex 5μ C18 100 Å column (250 × 4.6 mm) with a water:acetonitrile gradient containing 0.1% formic acid at 0.7 mL/min: 0-30 min, 5 to 50% acetonitrile. Using the Agilent MassHunter Quantitative Analysis software, extracted ion chromatograms were generated with 10 ppm mass windows around the calculated exact masses of protonated ormetoprim and desmethyltrimethoprim (*m/z* 275.1503 and *m/z* 277.1295, respectively). The areas of the peaks corresponding to the compounds were integrated and normalized to the dry weight of each sample.

### Statistical analysis

GraphPad Prism software Version 9.3.0 was used for statistical analysis. Two-tailed student’s unpaired t-tests were used to compare means between treatments (TMP or selected library compounds vs the vehicle solution DMSO), and between the efflux-deficient and -proficient strains treated with TMP. To compare means between strains treated with each of the library compounds and strains treated with DMSO, one-way ANOVA coupled with Dunnett’s multiple comparisons test were employed. Unpaired t-tests were also used to compare the relative abundance of compounds ormetoprim and desmethyltrimethoprim between efflux-proficient and efflux-deficient strains. Z’-factor calculation was done as described in Zhang *et al*. (58).

## Acknowledgments

This work was supported by NIH/NIAID R01 AI136794 (A.S.G & B.I.K.). We thank members of the Kazmierczak laboratory for their insightful comments on the manuscript and Stephen Lory for providing the PAO1 PA0397 (efflux-deficient) and isogenic wild-type control strains.

